# The algorithm for proven and young (APY) from a different perspective

**DOI:** 10.1101/2022.11.23.517757

**Authors:** Mohammad Ali Nilforooshan

## Abstract

The inverse of the genomic relationship matrix (**G**^−1^) is used in genomic BLUP (GBLUP) and the single-step GBLUP. The rapidly growing number of genotypes is a constraint for inverting **G**. The APY algorithm efficiently resolves this issue. Matrix **G** has a limited dimensionality. Dividing individuals into core and non-core, **G**^−1^ is approximated via the inverse partition of **G** for core individuals. The quality of the approximation depends on the core size and composition. The APY algorithm conditions genomic breeding values of the non-core individuals to those of the core individuals, leading to a diagonal block of **G**^−1^ for non-core individuals 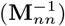. Dividing observations into two groups (*e.g*., core and non-core, genotyped and non-genotyped, *etc*), any symmetric matrix can be expressed in APY and APY-inverse expressions, equal to the matrix itself and its inverse, respectively. The change of **G**^*nn*^ to 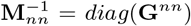 makes APY an approximate. This change is projected to the other blocks of **G**^−1^ as well. The application of APY is extendable to the inversion of any large symmetric matrix with a limited dimensionality at a lower computational cost. Furthermore, APY may improve the numerical condition of the matrix or the equation system.

## 1 Introduction

Genomic evaluations are mainly performed using the genomic relationship matrix **G** in the so-called method genomic BLUP (GBLUP, VanRaden (2008)) or random regression SNP marker models called SNP-BLUP (Koivula et al., 2012). The first predicts genomic breeding values of genotyped individuals, and the latter predicts marker effects (*i.e*., allele substitution effects). Simultaneous genetic evaluation of genotyped and non-genotyped individuals for obtaining optimal and unbiased evaluations not limited to genotyped individuals, both methods were elevated to single-step GBLUP (ssGBLUP, Aguilar et al. (2010); Christensen and Lund (2010)), and single-step SNP-BLUP (ss-SNP-BLUP, Fernando et al. (2014)), also called the single-step marker effect model.

The number of genotyped individuals is rapidly growing, and the most expensive operation in GBLUP and ssGBLUP is inverting matrix **G**. As the number of genotyped individuals reaches the number of markers, the numerical condition of **G** deteriorates. By the number of genotypes exceeding the number of markers, **G** becomes singular and non-invertible. Furthermore, the cost of inverting **G** and **A**_22_ (the block of **A** corresponding to genotyped individuals, where **A** is the pedigree-based additive genetic relationship matrix) required for ssGBLUP is cubic, and there is a bottleneck of direct inversion of a matrix of size about 150,000 (Fragomeni et al., 2015). Three solutions were proposed for this problem (Misztal et al., 2014; Fernando et al., 2016; Mäntysaari et al., 2017), one being the algorithm for proven and young (APY, Misztal et al. (2014)). This algorithm belongs to a group of methods called approximate kernel methods or Gaussian process approximations (Snelson and Ghahramani, 2007). APY forms a sparse representation Of 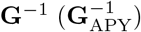, dividing genotyped individuals to core (*c*) and non-core (*n*) subsets. Direct inversion is only required for the block of **G** corresponding to core individuals (**G**_*cc*_). Consequently, the *O*((*c* + *n*)^3^) computational cost is reduced to *O*(*c*^3^) + *O*(*n*). In the APY algorithm, genomic breeding values of non-core individuals are conditioned on the genomic breeding values of core individuals. This algorithm is based on the assumption that the dimensionality of **G** is limited and that independent chromosome segments explain the rank of **G** (Misztal, 2016). As long as the number of core individuals is greater than the number of independent chromosome segments (Misztal et al., 2014), and the core subset covers the **G** spectrum (Bermann et al., 2022), it may not take all the genotyped individuals to explain the variation in **G**. Therefore, the variation in **G** can be explained by the core subset, and genomic breeding values of the non-core individuals are expressed as a linear function of those from the core individuals (Bermann et al., 2022). As such, the accuracy of the APY algorithm depends on the core size and composition.

The 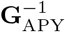 matrix is calculated as (Bermann et al., 2022):

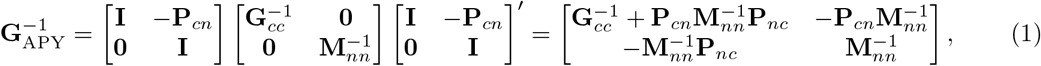

where, 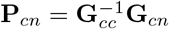 is a diagonal matrix with diagonal elements:

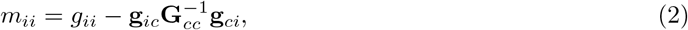

*g*_*ii*_ is a diagonal element of **G**_*nn*_, and **g**_*ic*_ is a row vector of **G**_*nc*_. Strandén et al. Strandén et al. (2017) and Bermann et al. Bermann et al. (2022) showed that:

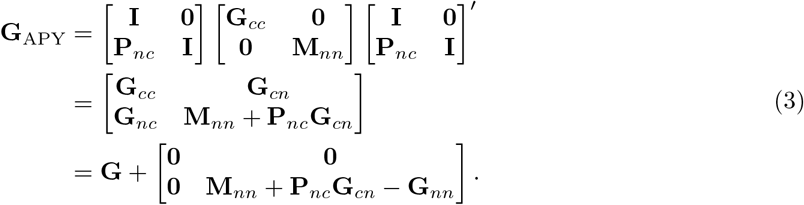

The aim of this study is to provide new insights and understanding about the APY algorithm.

## 2 Theory and Discussion

### 2.1 The APY and APY-inverse expressions

In this subsection, it is shown that any covariance or inverse covariance (generally any symmetric) matrix has expressions, here called APY and APY-inverse expressions. To understand the properties of the APY and APY-inverse expressions, we get help from the hybrid pedigree-genomic relationship matrix (**H**) used in ssGBLUP. Legarra et al. (Legarra et al., 2009) derived various forms of the same relationship matrix, including the full pedigree and genomic information. Denoting genotyped and non-genotyped individuals as 2 and 1: **H** =

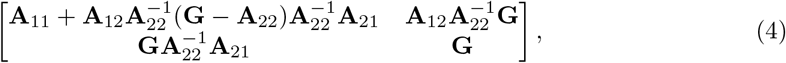

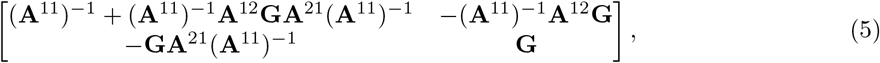

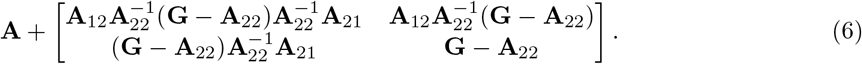

It worth mentioning that replacing **G** with **A**_22_ in any of these equations turns **H** to **A**. Similarly, replacing **G** with **G**_*nn*_ and **A** with **G** turns **H** to **G**. The above equations can be simplified to:

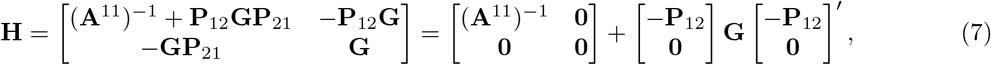

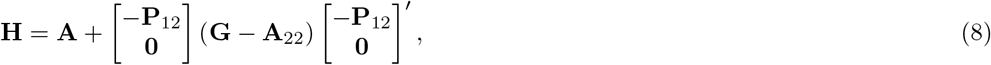

where, the projection matrix 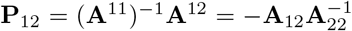. A nice property of **H** is that its inverse can be derived directly with no need to form and invert **H** (Aguilar et al., 2010; Christensen and Lund, 2010):

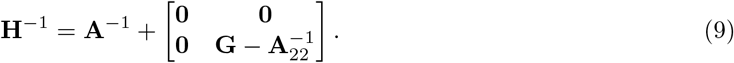

Matrix **H**^−1^ replaces **A**^−1^ in BLUP for ssGBLUP. Replacing **G** with 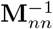, **A**^−1^ with **G**, and notations 1 and 2 with *c* and *n*, respectively, turns Eq. 7 to Eq. 1. This shows that Eq. 7 is the APY-inverse expression of **H**^−1^. Following Eq. 3, the APY expression of **H**^−1^ equals:

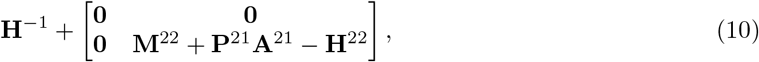

where **M**^22^ = **H**^22^ − **P**^21^**A**^12^, **P**^12^ = (**A**^11^)^−1^**A**^12^, and 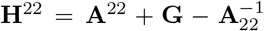. Similarly, there are APY and APY-inverse expressions for **H**. The APY and APY-inverse expressions of **G** are obtained by replacing 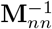 with **G**^*nn*^ in Eq. 3 and Eq. 1, respectively. Algebraically, it is possible to prove that the APY and APY-inverse expressions of **G** are equal to **G** and **G**^−1^, respectively.

### 2.2 Understanding the differences between G^−1^ and 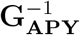

As shown in the previous subsection, replacing **G**^*nn*^ with 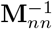 turns **G**^−1^ to 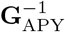. Note that, though, 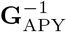 is an approximate **G**^−1^, the APY-inverse expression of **G** and the APY expression of **G**^−1^ are the exact **G**^−1^. In this subsection, the differences between **G**^*nn*^ and 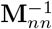, and between **G**^−1^ and 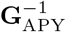 are explained.

Given Eq. 2, let’s reconstruct **M**_*nn*_ as if 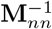 had non-zero off-diagonal elements to calculate.

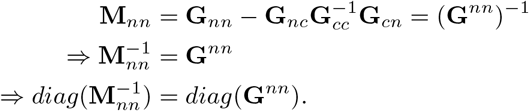

Thus, as long as the off-diagonal elements of 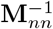 are not set to zero, 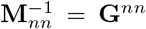. Eq. 2 is a computationally optimised way of calculating the reciprocals of the diagonal elements of **G**^*nn*^ (*i.e*., avoiding the calculations for off-diagonal elements, and the parallel computing possibility). Ignoring the off-diagonal elements of **G**^*nn*^ requires other block of **G**^−1^ compensate the missing information (making genomic relationships among non-core animals conditional to core animals), and this change should be projected to the other blocks of **G**^−1^ via the projection matrix **P**_*cn*_ (Eq. 1). Consequently, **G**^*nn*^ in the other blocks of **G**^−1^ are changed to 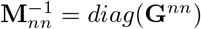to turn **G**^−1^ to 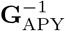 (Eq. 1).

In comparison with **G**, no change is made to **G**_APY_ other than to **G**_*nn*_ (Eq. 3). It can be articulated that the efficiency of the APY algorithm depends on how well **M**_*nn*_ + **P**_*nc*_**G**_*cn*_ represents **G**_*nn*_.

### 2.3 Other applications

The application of the APY algorithm is not limited to **G**^−1^, nor to GBLUP and ssGBLUP. This algorithm can be applied to approximate the inverse of any large symmetric matrix, where the rank of the matrix is smaller than its dimension. Representing any such matrix with **G**, only **G**_*cc*_ needs to be inverted. Besides reduced matrix inversion cost, there are sparsity-related reduced computational costs.

The first and the only time the APY algorithm was suggested for inverting a matrix other than **G** was by Misztal et al. Misztal et al. (2014). They suggested the APY algorithm for the **A**_22_ inversion, which is required in ssGBLUP (Eq. 9). They derived an equivalent formula for the APY approximation of 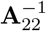:

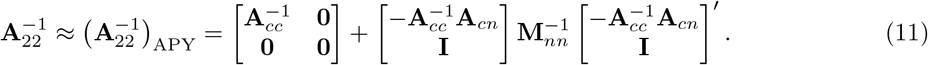

Here, the diagonal elements of **M**_*nn*_ equal 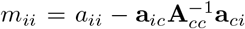, where *i* is a non-core genotyped individual. However, computing 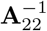 via the APY algorithm is a problem in a loop, which means to obtain the inverse of a block of **A** (*i.e*., 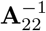), the inverse of its sub-block 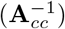 is required. There are two other well-established methods for the calculation of 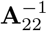 (Colleau, 2002; Faux and Gengler, 2013).

Contrarily, one may apply the APY algorithm for inverting **A**^−1^ to **A**. Though calculating **A** is computationally expensive, calculation of **A**^−1^ is computationally fast and efficient (Henderson, 1975), and it is sparse. The computational cost of inverting **A**^−1^ to **A** can be reduced by obtaining an APY inversion of **A**^−1^:

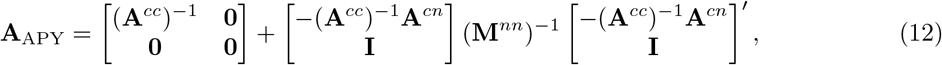

where **M**^*nn*^ is a diagonal matrix with diagonal elements *m*^*ii*^ = *a*^*ii*^ - **a**^*ic*^(**A**^*cc*^)^−1^**a**^*ic*^. If rather than **A**, a diagonal block of it (**A**_*cc*_) is needed, some of the calculations in Eq. 12 become redundant, and (**A**_*cc*_)_APY_ can be calculated as:

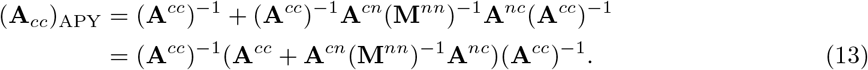

The above formula can be used to calculate an approximate **A**_22_ to blend with **G** (*i.e*., *k***G** + (1 − *k*)**A**_22_; 0 *< k <* 1) for improving the numerical condition of **G**, and to introduce residual polygenic variance not captured by the markers. However, the APY approximations of **A** and **A**_*cc*_ are expectedly inaccurate. For accurate approximations, non-core animals need to have their parents, progeny, and mates in the core group. The non-zero off-diagonal elements of **A**^−1^ are among parent-progeny and mates (Henderson, 1975). That considerably limits the choice and the number of animals in the non-core group. On the other hand, for 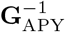, all animals in **G**_*cc*_ contribute information to each animal in **G**_*nn*_.

The APY algorithm helped overcome the limitations of inverting **G**. On the contrary, this constraint does not exist for marker effect models (*i.e*., SNP-BLUP and ss-SNP-BLUP) because a marker × marker matrix is used instead of **G**^−1^, which does not need to be inverted. This advantage comes at the price of dense matrix multiplications, and convergence complexities (Vandenplas et al., 2018; Bermann et al., 2022).

Unlike **G**, the size of that matrix remains constant over time unless the genotyping platform changes, and the old genotypes are imputed to a genotyping platform with a higher marker density. In fact, GBLUP and SNP-BLUP are equivalent models (Bermann et al., 2022). Conversion formulas between these two models are presented in the Appendix.

The mixed model equations (MME) of the marker effect models do not require direct matrix inversion (Fernando et al., 2014). Indirect inversion of **A** is needed, which is easy to obtain (Henderson, 1975). However, due to convergence difficulties, a specialised preconditioned conjugate gradient (PCG) solver with a block-diagonal Jacobi preconditioner matrix is applied, which is extended from single-trait to multi-trait analyses (Harris et al., 2022). As such, a marker × marker diagonal block of the MME (here called **Q**) is inverted, which is expanded by the number of traits in the model. The APY algorithm is a good candidate for this scenario, where the markers are divided into core and non-core. Only the block corresponding to core markers (**Q**_*cc*_) is inverted. Similar rules applied to 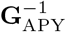 are applied to this scenario, with the difference that the role of markers and genotyped individuals are switched. Due to collinearity in the marker × individual genotype matrix, this matrix is not of full rank. The main source of collinearity is the markers with low minor allele frequency. Also, it would probably not take all the genotyped individuals to explain marker effects. Therefore, **Q** has a limited dimensionality, and the off-diagonal elements of **Q**_*nn*_ (in the preconditioner matrix, not in the MME) are conditioned on **Q**_*cc*_ and **Q**_*cn*_. The 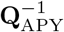 is a preconditioner matrix with the preconditioning properties similar to those of **Q**^−1^. Though the number of PCG iterations might differ, the cost of storing 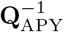 in the memory is cheaper, and each PCG iteration is expected to be faster due to the sparsity of 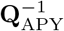

The core size and composition define the APY accuracy. Core size, which its optimum is a function of the effective population size (Pocrnic et al., 2016), is the most important. As long as there is room to increase the core size to span over 98% of the eigenvalue spectra of **G**, a random set of core individuals is shown to perform well because it gives good coverage over generations and breeds in the population. With a sufficiently large core size, the gain from an optimal core subset is marginal (Nilforooshan and Lee, 2019). There is ongoing research on finding the optimal core subset, and it is an important topic for admix populations and when the core size is constrained. When the core size is limited, an optimum core composition can harvest a larger variation of **G**, and provide more accurate estimated of breeding values (Abdollahi-Arpanahi et al., 2022; Pocrnic et al., 2022).

It is unknown what proportion of random markers would cover over 98% of the eigenvalue spectra of **Q**. Similar to the concept of effective population size defining the optimum number of core individuals for 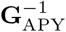 for 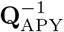 might be the concept of effective marker size defining the optimum number of core markers. Such markers are likely segregating in the coding regions, with effects as independent and orthogonal as possible to other markers; a concept similar to independent chromosome segments equal to 2*N*_*e*_*L/log*(4*N*_*e*_*L*) (Goddard, 2009), where *N*_*e*_ is the effective population size, and *L* is the length of chromosome in Morgans. Therefore, **G** and **Q** might have similar dimensionality, and the required core size might be the same for both. Possibly, choosing markers corresponding to the highest diagonal elements of **Q** is better than a random set of core markers. This is because those markers cover a larger variation in **Q** (*i.e*., *trace*(**Q**) = Σ*eigenvalue*(**Q**)). This would favour choosing markers with lower minor allele frequency. An optimised core subset may reduce the need for a larger core size (*i.e*., the same variation in **Q** captured by a smaller set of markers). Future research is needed on this topic.

The APY accuracy is usually measured by the correlation between genomic breeding values obtained via **G**^−1^ and 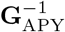. However, it might be okay to have a correlation coefficient slightly less than 1. A small variation of **G** might be due to collinearity and noise-related and good to get discharged. The APY algorithm may help reduce the collinearity and noise in **G**. Nilforooshan and Lee Nilforooshan and Lee (2019) showed that APY reduced the very large max(diag(**G**^−1^)), which is a sign of reduced collinearity and improved condition of **G**. Validation of genomic breeding values is a good complementary.

Another benefit of APY is reducing the computational cost of blending **G** with **A**_22_ (Hollifield et al., 2022). APY does not require storing **G**_*nn*_, but *diag*(**G**_*nn*_). Therefore, rather than calculating the whole **A**_*nn*_, calculation of *diag*(**A**_*nn*_) = 1 + **F**_*n*_ would be enough, where **F**_*n*_ is the vector of inbreeding coefficients for *n*. The inbreeding coefficients, however, need to be calculated before calculating any row of **A**_22_ (Colleau, 2002), including the rows for *c*. The average of the diagonal and off-diagonal elements of **A**_22_ might be needed for tuning **G**. However, it can be assumed that *µ*(*offdiag*(**A**_*nn*_)) = 0 (*i.e*., *µ*(*offdiag*(**A**_*cc*_)) + *µ*(**A**_*cn*_) = *µ*(*offdiag*(**A**_22_))).

## 3 Conclusions

This study aimed to open new insights and understanding about the APY algorithm. It was shown that any symmetric matrix can be expressed as a combination of its two diagonal blocks (for core and non-core animals, genotyped and non-genotyped animals, *etc*). The projection matrix makes the combination (information flow) between the two diagonal blocks. As such, any symmetric matrix has (the so-called) APY and APY-inverse expressions equal to the matrix itself and its inverse, respectively. The difference arises when the off-diagonal elements of a diagonal block of the APY-inverse expression (corresponding to the non-core set) are set to 0. This change is projected to the rest of the inverse matrix via the projection matrix. That diagonal matrix sets non-core individuals independent from each other conditional to the coefficients provided by the core individuals. The APY algorithm can also be understood as an approximate absorption of the off-diagonal elements of a diagonal block into the rest of the matrix.

The APY algorithm is based on the concept of the limited matrix dimensionality. A genomic relationship matrix has a limited dimensionality equivalent to the number of independent chromosome segments, which allows a reduction in the dimensionality of **G**. Therefore, it would take the inverse of a diagonal block of **G** to invert **G**. An APY-inverse of **G** with a sufficient core size and proper core composition produces genomic breeding values analogous to those using the exact **G**^−1^. A possible new application for APY is computing the block corresponding to marker effects, of the block-diagonal preconditioner matrix for iterative solving of marker effect model equations. The application of APY is not limited to obtaining the best sparse approximates of **G**^−1^, and new applications may emerge in the future.

## Appendix

Considering the MME for GBLUP:

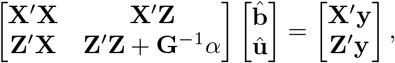

and **G** = **WW**^*′*^, conversion of GBLUP to SNP-BLUP MME follows:

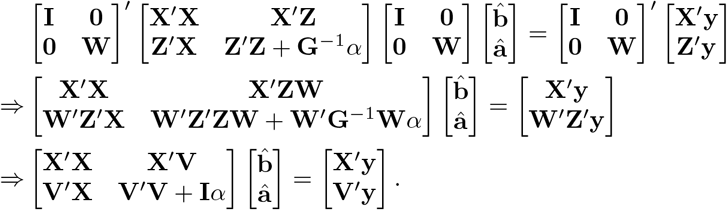

On the other hand, the conversion of SNP-BLUP to GBLUP is as follows:

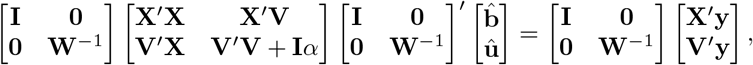

where 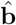, **û** and **â** are the vectors of solutions for fixed effects, individuals’ additive genetic merit and marker effects, 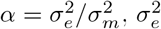 is the residual variance, and 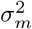 is the additive genetic variance captured by markers.

